# Molecular mechanisms of substrate-controlled ring dynamics and sub-stepping in a nucleic-acid dependent hexameric motor

**DOI:** 10.1101/069559

**Authors:** Nathan D. Thomsen, Michael R. Lawson, Lea B. Witkowsky, Song Qu, James M. Berger

## Abstract

Ring-shaped hexameric helicases and translocases support essential DNA, RNA, and protein-dependent transactions in all cells and many viruses. How such systems coordinate ATPase activity between multiple subunits to power conformational changes that drive the engagement and movement of client substrates is a fundamental question. Using the *E. coli* Rho transcription termination factor as a model system, we have employed solution and crystallographic structural methods to delineate the range of conformational changes that accompany distinct substrate and nucleotide cofactor binding events. SAXS data show that Rho preferentially adopts an open-ring state in solution, and that RNA and ATP are both required to cooperatively promote ring closure. Multiple closed-ring structures with different RNA substrates and nucleotide occupancies capture distinct catalytic intermediates accessed during translocation. Our data reveal how RNA-induced ring closure templates a sequential ATP-hydrolysis mechanism, provide a molecular rationale for how the Rho ATPase domains distinguishes between distinct RNA sequences, and establish the first structural snapshots of substepping events in a hexameric helicase/translocase.

**SIGNIFICANCE:** Hexameric, ring-shaped translocases are molecular motors that convert the chemical energy of ATP hydrolysis into the physical movement of protein and nucleic acid substrates. Structural studies of several distinct hexameric translocases have provided insights into how substrates are loaded and translocated; however, the range of structural changes required for coupling ATP turnover to a full cycle of substrate loading and translocation has not been visualized for any one system. Here, we combine low-and high-resolution structural studies of the Rho helicase, defining for the first time the ensemble of conformational transitions required both for substrate loading in solution and for substrate movement by a processive hexameric translocase.

## INTRODUCTION

Hexameric helicases and translocases are motor proteins that play a central role in cellular transactions ranging from replication and repair to transcriptional regulation, chromosome packaging, and proteolytic homeostasis (1-4). Used to drive the processive and at times highly-rapid movement of extended nucleic acid or protein chains through a central pore, ring-shaped motors face several challenges to their operation. One is that certain enzymes must transition through controlled ring opening and/or subunit assembly events to allow long, polymeric substrates that lack freely-accessible ends access to interior motor elements (5-7). Another is that, once loaded, the molecular plasticity inherent to these assemblies must be harnessed to precisely coordinate ATP binding and hydrolysis between multiple subunits for powering substrate translocation, while at the same time alternating between tight and loose grips on the substrate to allow for processive movement. The substrate-dependent molecular rearrangements that underpin ring dynamics during these events remain poorly understood not only for hexameric helicases and translocases, but for related ring-shaped switches as well.

The *E. coli* Rho transcription termination factor is a well-established model system for understanding hexameric translocase and helicase function (8, 9). During termination, Rho uses a cytosine-specific RNA binding domain appended to the N-terminus of a RecA-type ATPase fold (10, 11) to bind nascent RNA transcripts at cytosine-rich sequences (known as Rho utilization (*rut*) sites) (12, 13). Once loaded, Rho consumes ATP to translocate 5’→3’ toward a paused RNA polymerase, eventually promoting transcription bubble collapse and RNA release (14-19).

Structural studies have provided insights into the mechanics of RNA loading by Rho, imaging the helicase in both closed-circular and notched/lockwasher-shaped states (20-24). These findings have suggested that Rho monomers assemble into a pliant hexamer that can spontaneously open to allow nucleic acid entry into the translocation pore (22, 24, 25), a property that has been observed in other hexameric helicases such as the eukaryotic MCM2-7 complex (26-28). By comparison, the structure of a closed-ring Rho complex bound to single-stranded RNA and the ATP-mimetic ADP•BeF_3_ has provided a picture of a prospective translocation intermediate of the motor (29). This work, combined with that of a prior landmark model for the human papillomavirus E1 helicase bound to single-stranded DNA and ADP (30) (as well as more recent work on the replicative helicase, DnaB (31)), has led to two general principles for understanding hexameric helicase function. One is that, as proposed first for the T7 gp4 protein (32), hexameric helicases likely employ a sequential, rotary ATPase mechanism similar to that used by the F1 ATPase to power translocation (33). Another is that both the relative order of hydrolysis between subunits and/or differences in subunit orientation within a motor can determine the polarity of nucleic acid translocation (29, 30, 34, 35).

Despite present-day insights, there remain many unanswered questions regarding the loading and translocation mechanism of Rho. For example, there has been no systematic assessment of the extent to which the various crystallographic states observed thus far correlate with the predominant structures adopted by Rho in solution. The relative roles that ATP and RNA play in promoting ring opening *vs*. ring closing are similarly unresolved. At the same time, the range and types of movements that accompany ATP turnover events to support RNA translocation have yet to be fully defined. Likewise, it is unclear how the physical nature of the nucleic acid substrate itself – its chemical identity and base composition – feeds back to ATPase centers to influence motor activity. Answers to these questions are needed to understand fundamental aspects of hexameric helicase loading and translocation not just by Rho, but by hexameric motor proteins in general.

To better define the physical basis by which substrate-coupled conformational changes underpin hexameric motor mechanism, we conducted both solution and crystallographic structural studies of *E. coli* Rho with the goal of characterizing novel conformational intermediates in conjunction with different RNA sequences and nucleotide-binding modes. Small-angle X-ray scattering (SAXS) confirms that the current open- and closed-ring models imaged crystallographically correspond to the dominant states adopted in solution in the absence and presence of substrates, respectively, and further establish that Rho ring closure occurs only when both RNA and ATP are present. Three new crystal structures of Rho (including an experimentally-phased structure at higher resolution than that determined previously) reveal new ring rearrangements accompanying ATP cycling, which together with comparisons to other oligomeric RecA-family motors, uncover a common allosteric linkage that physically connects nucleic-acid binding with ATPase activation in members of this widespread motor family.

Together, our data highlight unexpected sub-steps in the Rho loading/translocation cycle, and help explain previous research implicating RNA sequence as a regulator of Rho function in transcription termination (36-39).

## RESULTS

### Both RNA and ATP are required to drive Rho ring closure

To probe how Rho physically and differentially responds to the binding of substrates such as nucleotide and RNA, we first used small angle X-ray scattering (SAXS) to verify whether the available crystal structures of the helicase correspond to major or minor population states in solution (**Fig. 1a-b and Supplementary Fig. 1a-b**) (40). SAXS data were initially collected on Rho in the absence of ligand, and on Rho mixed with one molar equivalent per hexamer of a poly-pyrimidine RNA (rU12) and 2 mM ADP•BeF3, a non-hydrolyzable ATP analog. Consistent with size-exclusion chromatography (**Supplementary Fig. 1c-d**), SAXS traces for Rho exhibited no sign of aggregation under any of the conditions tested, and distance distribution (P_(r)_) profiles, Guinier analysis, and Kratky plots also were consistent with a well-behaved, hexameric particle (**Supplementary Fig. 2 and Supplementary Table 1**). Notably, data collected in the absence and presence of both RNA and ADP•BeF3 proved to be extremely close matches to the theoretical curves calculated from models of the open and closed-ring structures of Rho, respectively (**Fig. 1a-b**). These data indicate that the open and closed-ring crystal structures seen previously for Rho indeed reflect the predominant conformations of native particles in solution.

**Figure 1.**
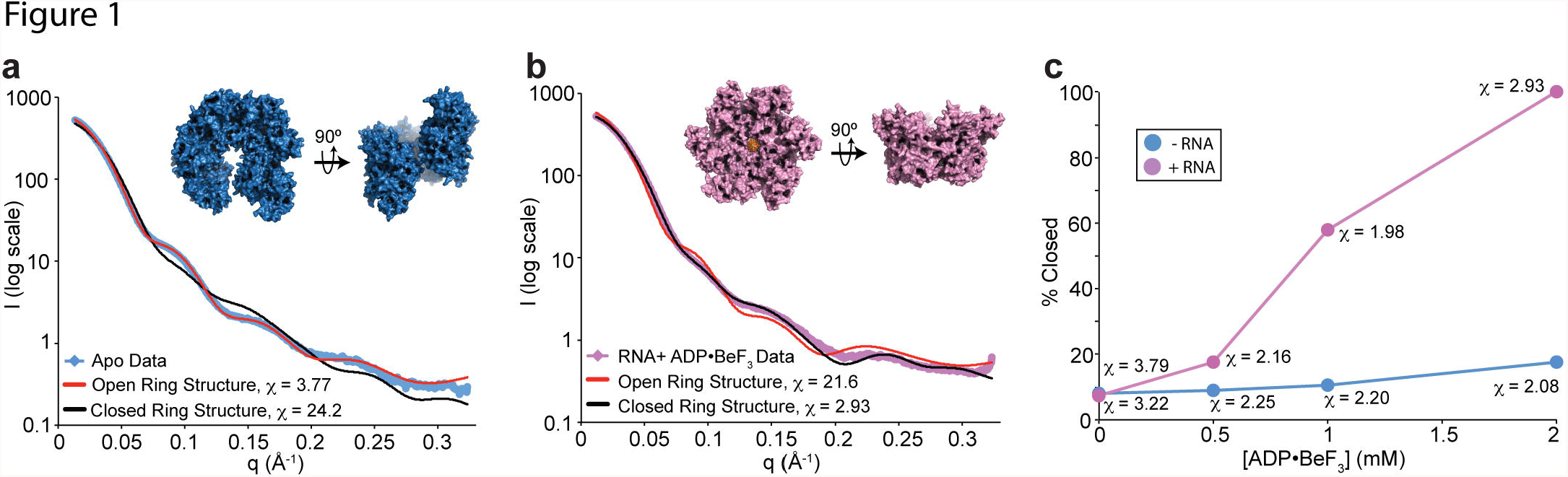
SAXS analysis of apo and substrate-bound Rho complexes. (a) Scattering curve for apo Rho in solution (blue) closely matches the theoretical curve (red) calculated from a model based on the open-ring structure (1PVO, inset (22)), but not the theoretical curve (black) calculated from a closed ring-Rho model bound to RNA and ADP•BeF_3_. (b) Scattering curve for Rho in the presence of RNA (1:1 RNA:hexamer) and 2 mM ADP•BeF_3_ (pink) closely matches the theoretical curve (black) calculated from the closed ring-Rho structure bound to RNA and ADP•BeF_3_ determined here but not the theoretical curve (red) calculated from a open ring Rho model (1PVO). (c) Plot of the percent-closed value obtained with FOXS *versus* ADP•BeF _3_concentration shows that both RNA and ATP are required for Rho ring closure.

To assess the individual contributions of RNA and nucleotide to ring closure, we next conducted SAXS experiments in either the presence or absence of RNA, and over several ADP•BeF3 concentrations. Data collected in the absence of RNA but with 0, 0.5, 1, or 2 mM ADP•BeF3 induced only modest changes in the scattering profile (**Supplementary Fig. 1a**). By contrast, scattering data collected in the presence of RNA exhibited distinct changes as ADP•BeF3 concentrations were increased (**Supplementary Fig. 1b**). Data collected at intermediate concentrations of nucleotide could not be properly modeled by individual Rho crystal structures, so we used the programs FOXS and OLIGOMER to fit a two-component model consisting of both the open- and closed-ring Rho structures (**Supplementary Tables 2-3**) (41-43). The two-component analysis produced greatly improved fits, indicating that in the absence of RNA, ∼92% of the Rho hexamers are open in solution, and that ∼82% remain open even at high (2mM) ADP•BeF_3_ concentrations (**Fig. 1c**). By contrast, with the addition of RNA, nucleotide titration leads to ring closing in Rho, such that by 2mM ADP•BeF_3_, the hexamer population approaches 100% closed. Together, these data confirm that Rho forms an open lock-washer in solution, and further demonstrate that both RNA and nucleotide are required to promote ring closure (5). This RNA- and ATP-dependent conformational change in Rho may be the rate-limiting step previously observed in stopped-flow kinetics measurements of ATP hydrolysis (44), a notion that is supported by the noted acceleration of both ATPase and ring closure rates in response to primary site occupancy ((38) and (45)).

### A new closed-ring Rho state manifests a sequential shift in ATPase status

Having established that existing crystallographic models are suitable representations of predominant Rho states in solution, we next set out to determine whether different substrate-bound, closed-ring forms of Rho might exist. We postulated that different RNA sequences or nucleotide-binding conditions might influence the formation of such intermediates. Since our initial closed-ring Rho crystals were difficult to work with (hereafter referred to as Rho^PolyU-P1^, in reference to the RNA sequence and space group in which the protein crystallized) (46), we set out to find a new crystal packing arrangement that could be obtained in the presence of different RNA and nucleotide conditions, but that still possessed at least one complete Rho hexamer per asymmetric unit (so as to image the particle in the absence of crystal symmetry influences). We therefore conducted a new, broader set of screens with purified Rho in the presence of RNA and nucleotide, discovering a crystal form belonging to the spacegroup P2_1_ that satisfied these criteria (**Methods**).

From the new Rho form, we were able to experimentally phase and refine a higher resolution (2.6 Å) structure of an intact hexamer bound to seven nucleotides of a centrally bound poly-U RNA, and that displayed electron density for ADP•BeF_3_•Mg^2+^ bound to all six ATP-binding sites (hereafter referred to as Rho^PolyU-P2_1_^) (**Fig. 2a-d and Table 1**). The experimental phases and higher resolution allowed us to correct two small building errors in previous Rho structures (**Supplementary Fig. 3**). A global alignment of the new P2_1_ model with the original P1 state reveals that the two hexamers are highly similar structurally (**Fig. 2a**), with a root mean squared deviation (RMSD) for all shared backbone Cα atoms of 1.55 Å. As seen previously, five subunits of the Rho^PolyU-P2_1_^ state form a smoothly-ascending spiral staircase that wraps around a single-stranded RNA helix; the sixth (subunit F) occupies a “transit” position midway between the uppermost (A) and bottommost (E) protomers to close the ring (**Fig. 2b**). In addition, four classes of nucleotide-binding sites were again evident at the six intersubunit interfaces. Using criteria such as nucleotide B-factors (a measure of positional accuracy, or “orderedness” (47)) and coordination geometry, these sites have been classified as corresponding to either “exchange” (*E*), “ATP-bound” (*T*), “hydrolysis” (*T^*^*), or product (*D*) states along the ATPase cycle (**Fig. 2a**) (29).

**Figure 2.**
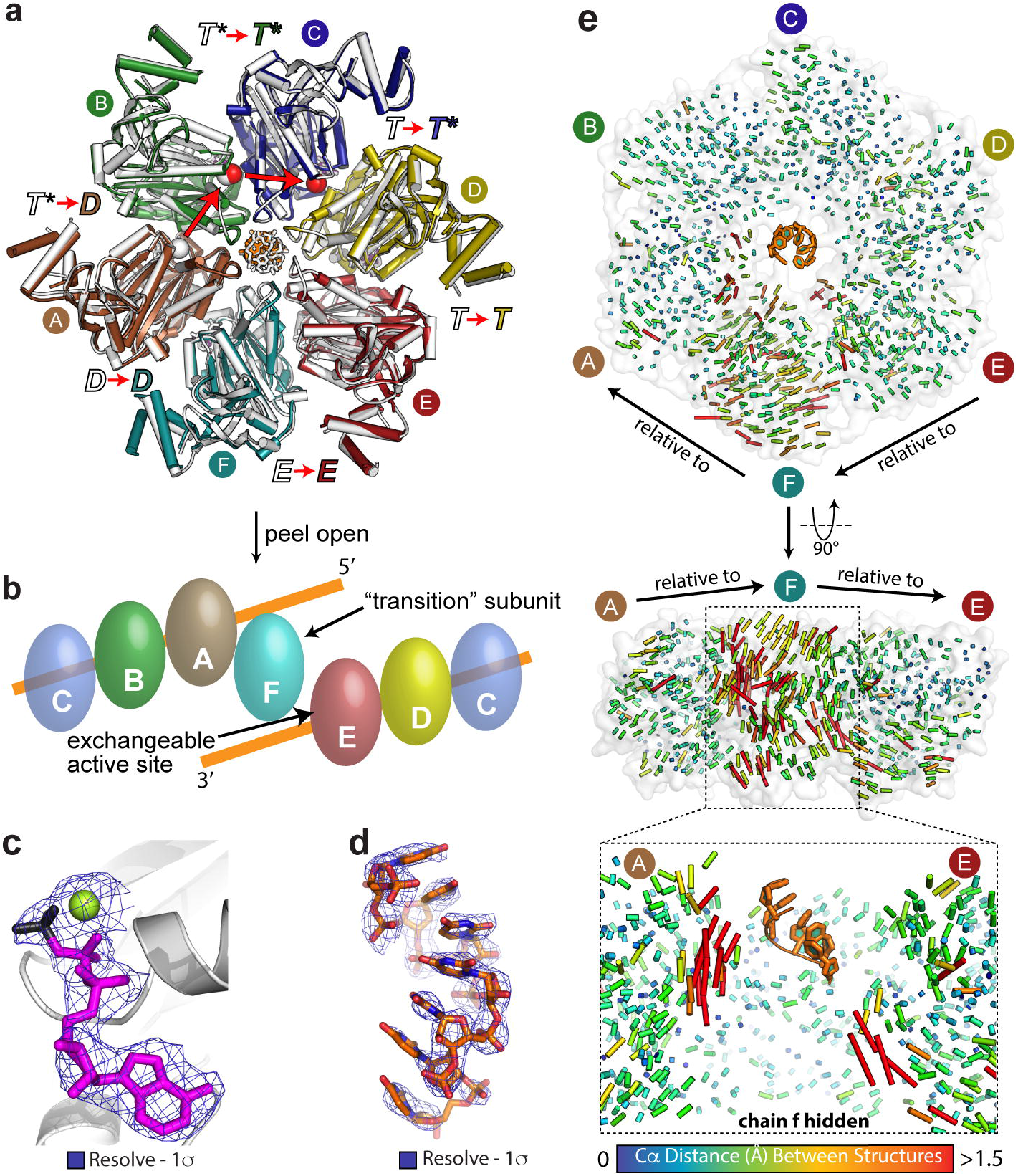
Overall structure and ligand binding in a new Rho translocation intermediate. (**a**) Superposition of a new Rho•RNA•ADP•BeF3 structure (Rho^PolyU-P2_1_^, colored and labeled by subunit), with the previously-determined Rho•RNA•ADP•BeF3 structure (Rho^PolyU-P1^, white). Presumptive ATP hydrolysis states ( *T*– ATP bound, *T^*^*– ATP hydrolysis, *D*– product, *E*– exchange) are labeled in color (Rho^PolyU-P1^) or white (Rho^PolyU-P1^). (**b**) Schematic of closed-ring Rho hexamer (flattened on page) bound to RNA (one complete turn) highlighting the right-handed helical rise of the A/B/C/D/E interfaces that are bound to RNA, and the position of subunit F in transition between the top and bottom steps of the RNA binding staircase. Subunit C is shown twice to connect the two sides. (**c**) Experimental (density modified, SAD phased, Rho^PolyU-P2_1_^) electron density around one of the bound nucleotides (ADP – magenta, BeF _3_– black, Mg^2+^ – yellow-green). (**d**) Experimental (density modified, SAD phased, Rho^PolyU-P2_1_^) electron density around the bound RNA (orange). (**e**) Intersubunit distance vectors calculated independently for each intersubunit interface between the Rho^PolyU-P1^ and Rho^PolyU-P2_1_^ structures. Vectors were calculated relative to either the clockwise-adjacent subunit (top) or to the counterclockwise-adjacent subunit (bottom) revealing that the new P2_1_ structures differ from the original P1 model primarily in the position of subunit F, and in the Q and R-loops of subunits A and E.

**Table 1.** Data collection, phasing and refinement.

|  | Rho<sup>PolyU-P2<sub>1</sub></sup> | Rho<sup>PolyU-P2<sub>1</sub></sup> | Rho<sup>PolyA</sup> | Rho<sup>Empty</sup> |
| --- | --- | --- | --- | --- |
| **Data collection** |  |  |  |  |
| Space group | P2<sub>1</sub> | P2<sub>1</sub> | P2<sub>1</sub> | P2<sub>1</sub> |
| Cell dimensions |  |  |  |  |
| *a, b, c* (Å) | 68.72, 198.8, 111.8 | 69.00, 198.6, 111.4 | 68.76, 199.2, 112.2 | 69.09, 199.1, 111.7 |
| $\alpha, \beta, \gamma$ (°) | 90.0, 104.8, 90.0 | 90.0, 104.4, 90.0 | 90.0, 104.8, 90.0 | 90.0, 105.0, 90.0 |
|  | *peak* | *remote* | *remote* | *remote* |
| Wavelength | 0.9796 | 1.116 | 1.116 | 1.116 |
| Resolution (Å) | 50-2.95 (3.06-2.95) | 50-2.60 (2.69-2.60) | 50-3.15 (3.26-3.15) | 50-3.2 (3.31-3.20) |
| *R*<sub>sym</sub> or *R*<sub>merge</sub> | 9.1 (46) | 8.1 (63) | 12.1 (60) | 12.8 (58) |
| *I*/σ*I* | 10.8 (2.4) | 14 (1.8) | 9.4 (2.0) | 8.1 (1.8) |
| Completeness (%) | 98.2 (97.1) | 99.7 (99.1) | 99.0 (99.5) | 96.2 (96.4) |
| Redundancy | 5.6 (5.2) | 3.1 (3.1) | 2.9 (2.9) | 2.8 (2.8) |
| **Refinement** |  |  |  |  |
| Resolution (Å) |  | 47.4-2.6 | 49.8-3.15 | 49.5-3.2 |
| No. reflections |  | 88,380 | 49,848 | 46,060 |
| *R*<sub>work</sub>/*R*<sub>free</sub> |  | 21.8 / 24.6 | 23.3 / 26.5 | 23.2 / 26.6 |
| No. atoms |  |  |  |  |
| Protein |  | 19,164 | 18,696 | 18,888 |
| Ligand/ion |  | 341 | 347 | 325 |
| Water |  | 271 | 19 | 15<sup>a</sup> |
| B-factors |  |  |  |  |
| Protein |  | 67.9 | 80.7 | 73.7 |
| Ligand/ion |  | 53.6 | 69.9 | 66.6 |
| Water |  | 45.2 | 64.5 | 46.6 |
| R.M.S deviations |  |  |  |  |
| Bond lengths (Å) |  | 0.006 | 0.006 | 0.006 |
| Bond angles (°) |  | 0.575 | 0.560 | 0.691 |

Values in parenthesis correspond to the highest resolution bin
<sup>a</sup>Only waters involved in octahedral coordination of the active site Mg<sup>2+</sup> ions were modeled

Globally, the conformations of the ATPase sites and the sequential order of ATPase states in the new P2_1_ Rho model initially appeared similar to those seen in the original P1 structure, with the modeled nucleotides refining to a relative B-factor distribution consistent with the formation of three tight and three weak ATP-binding environments as observed biochemically (48-53) (**Supplementary Fig. 4**). However, inspection of the distance vectors calculated between Cα atoms for the P1 and P2_1_ models revealed that subunit F, as well as the RNA binding R-loops of subunits E and A, have moved with respect to one other around a vector roughly parallel to the ring axis (**Figure 2e**).

Because the A, F and E subunits are thought to represent the likely sites for the release of nucleotide hydrolysis products and the binding of fresh ATP within the hexamer (29), we elected to compare the nucleotide-binding regions of the two Rho models in more detail. This analysis uncovered pronounced changes in some of the nucleotide-binding sites, which in turn revealed a shift in the pattern of functional states between the P1 and P2_1_ forms of Rho (**Figs. 2a, 3**). The most striking change is that the presumptive catalytic water molecules seen in subunit A and B active sites of the initial closed ring structure now appear in the catalytic centers of subunits B and C in the new structure (**Fig. 2a, 3a-c and Supplementary Fig. 5a-c**). Moreover, each of these waters also has shifted its local position from a non-ideal location in the Rho^PolyU-P1^ model to now take up a direct, in-line attack configuration with the BeF3 group of the bound nucleotide in the Rho^PolyU-P2_1_^ structure (**Fig. 3b**). Repositioning of the waters is mediated at both interfaces by a conserved arginine (the “arginine-switch” (RS), R269), the catalytic glutamate (CE) residue (E211) that polarizes the water to assist hydrolysis, and a backbone carbonyl (G337) of the adjacent protomer. The glycine-mediated contact was not observed in the prior P1 Rho structure. Consistent with biochemical studies implicating their importance to overall Rho function (10, 54-58), a conserved network of ion-pairs at the B/C and C/D interfaces – which runs from the RNA-binding pore to the ATPase center – appears to help orient amino acids responsible for in-line placement of the attacking water (**Supplementary Fig. 6**). Overall, the new placement for the catalytic waters in Rho^PolyU-P2_1_^, together with other changes in the relative orientations of catalytic important amino acids (**Fig. 3a-f and Supplementary Discussion**), supports the proposed existence of hydrolysis-competent active sites in the closed-ring Rho states (29). This shift also advances the relative position of the two *T^*^*states by one subunit toward the lone *E*-state, and adds a second *D*-state compared to the Rho^PolyU-P1^ structure determined previously (**Fig. 2a**).

**Figure 3.**
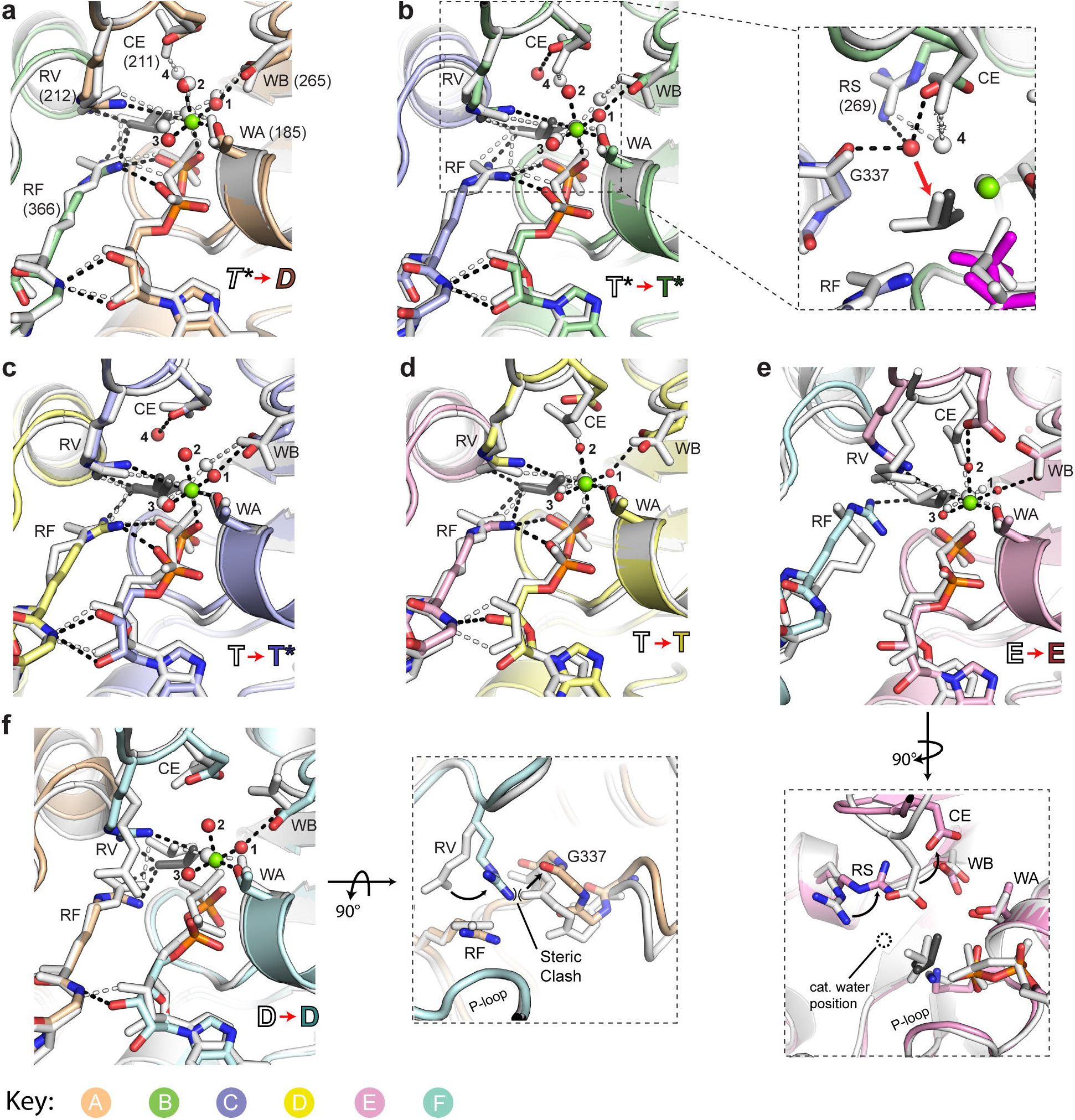
Changes in ADP•BeF_3_ coordination as revealed by superpositions between six Rho^PolyU-P1^ (white) and Rho^PolyU-P2_1_^ (colored) ATPase centers. In all subunits, changes are consistent with a partial shift in catalytic status of the six active sites around the ring. Abbreviations: WA – Walker A, WB – Walker B, RS – arginine switch, RF – arginine finger, RV – arginine valve. (**a**) Subunit A/B interface. While a water molecule is observed to associate with the catalytic glutamate (CE) at this site in Rho^PolyU-P1^ (water 4), it is not observed in the new Rho^PolyU-P2_1_^ structure. (**b**) Subunit B/C interface. Both catalytic centers show bonding interactions (dashed lines) characteristic of the *T* and *T^*^* states. Inset shows a zoomed-in view of the catalytic center with the RV removed (for clarity): whereas the prospective attacking water molecule in Rho^PolyU-P1^ was not ideally positioned for in-line chemistry with the γ-phosphate moiety (mimicked by the BeF3 – white), the CE-associated water molecules seen in the *T^*^* states of the new Rho^PolyU-P2_1_^ are positioned for ideal in-line attack (red arrow). (**c**) Subunit C/D interface. Concomitant with the disappearance of the catalytic water molecule in the A/B subunit interface of the Rho^PolyU-P2_1_^ structure, a catalytic water molecule appears in the C/D subunit interface compared to the Rho^PolyU-P1^ structure. (**d**) Subunit D/E interface. This site retains contacts characteristic of the *T* state in both structures. The CE in Rho^PolyU-P2_1_^ has broken its atypical contact with the Mg2% ion observed in Rho^PolyU-P1^, suggesting that it is transitioning to a slightly more active arrangement. (**e**) Subunit E/F interface. The arginine finger in the Rho^PolyU-P1^*E*-state appears to be pointing out of the active site, while the arginine finger in Rho^PolyU-P2_1_^ clearly points into the active site and coordinates the bound BeF3. However, the arginine valve in the Rho^PolyU-P2_1_^ structure is pulled slightly out of the active site, resulting in the same number of contacts between the arginine groups and the BeF3 molecule. The catalytic glutamate in this active site is in an atypical conformation in both structures, interacting with the Mg^2%^ ion directly (P1) or indirectly *via* coordinated waters (P21). Rotated view – the catalytic glutamate and arginine switch in Rho^PolyU-P2_1_^ are moved out of the catalytic site in the *E*-state when compared to the Rho^PolyU-P1^ structure. (**f**) Subunit F/A interface. The arginine valve in the Rho^PolyU-P1^*D*-state points out of the active site, while the arginine valve in the Rho^PolyU-P2_1_^ structure points into the active site and coordinates the bound BeF3. Rotated view – the arginine valve now points into the active site due to an opening of the intersubunit interface in Rho^PolyU-P2_1_^ relative to Rho^PolyU-P1^ that relieves a steric clash between the guanidinium group of the arginine and G337 of the adjacent subunit.

### Inter-subunit relationships in the nucleotide-exchange region mirror those of an open-ring state

A key supposition of the rotary cycling model based on the polyU-bound P1 Rho crystal form has been that the E/F interface, which is relatively open and solvent exposed, corresponds to a nucleotide exchangable (*E*) site where substrate can bind to or product can dissociate from the hexamer, respectively (29). To test this idea, we back-soaked nucleotide out of the new Rho^PolyU-P2_1_^ crystal form and determined the structure of the helicase by X-ray crystallography. Inspection of all active sites in the model revealed very weak and discontinuous electron density for the nucleotide-binding site of subunit E, indicating a very low occupancy for ADP•BeF3 that precluded modeling of nucleotide into the site (nucleotide density in the other active sites was unchanged) (**Fig. 4a-b**). Intersubunit distance vectors calculated between the back-soaked structure (referred to hereafter as Rho^Empty^) and the fully occupied Rho^PolyU-P2_1_^ structure show that the relative positions of subunits E and F shift by ∼1.7 Å with respect to one another, but that the rest of the subunits in the hexamer undergo only minor positional changes (**Fig. 4c**). A comparison of the E/F interface in the presence and absence of ADP•BeF3 reveals an increased separation of the two subunits that moves the Arg-finger out of the active site and that buries ∼120Å^2^ less total surface area at the interface (**Fig. 4d**). Together, these observations further confirm the proposition that the E/F active site corresponds to a nucleotide-exchange point, and show that product release coincides with an increased opening of the subunit interface at this catalytic center. Interestingly, upon the disappearance of nucleotide from the *E*-state, the relative B-factors for the nucleotide in the active site of subunit F also increased relative to other Rho structures (**Supplementary Fig. 4**), suggesting that product release may cooperatively weaken the association of nucleotide in the immediately adjacent *D*-state.

**Figure 4.**
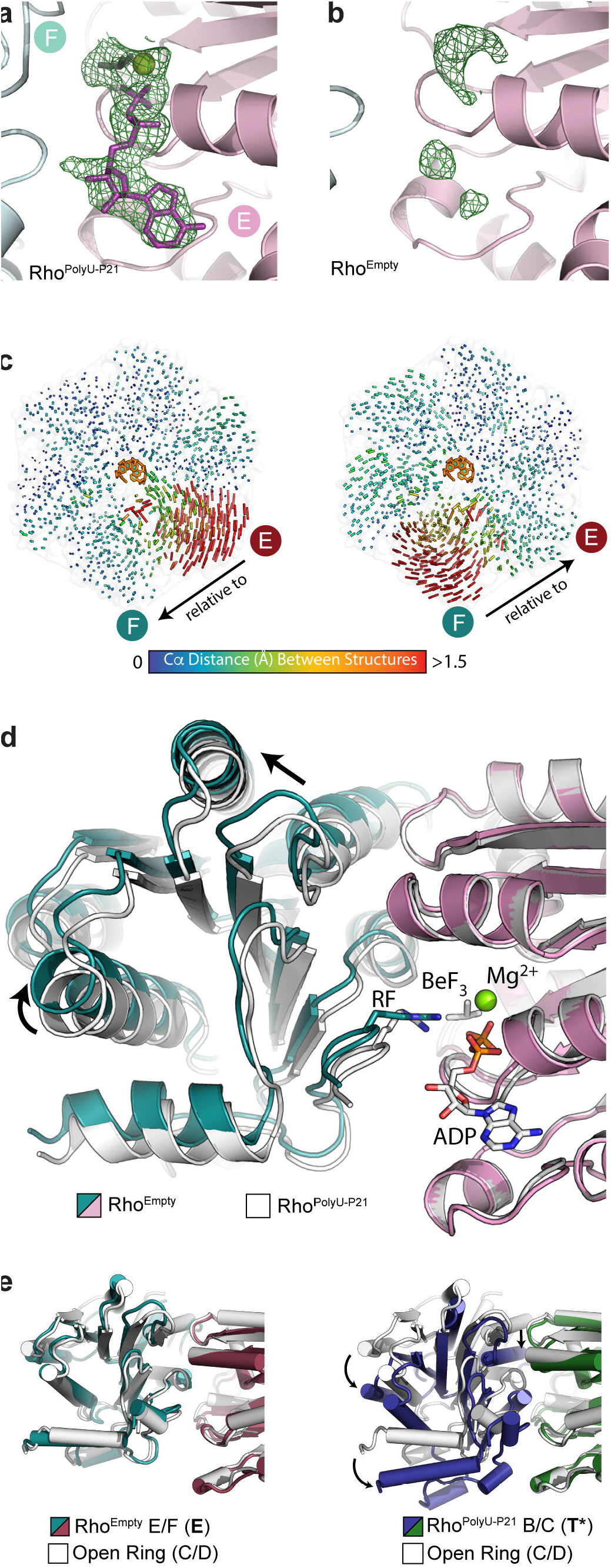
Opening of the active site in the exchange (*E*) state (subunit E) accompanies nucleotide release. (**a**) Simulated annealing omit map of the *E*-state in Rho^PolyU-P2_1_^ revealing strong density for the bound nucleotide. (**b**) Simulated annealing omit map of the *E*-state in the Rho^Empty^ structure reveals poor density suggesting low nucleotide occupancy and/or order. (**c**) Intersubunit distance vectors calculated independently for each intersubunit interface and comparing RhoEmpty and Rho^PolyU-P2_1_^ structures. The vectors reveal that the only significant intersubunit changes between these structures are at the E/F interface. (**d**) Superposition of chain E from the Rho^PolyU-P2_1_^ and RhoEmpty structures reveals that nucleotide release induces a rigid body shift in chain F (arrows) that causes the active site to open. The arginine finger (RF), which also changes position, is labeled. (**e**) Structural alignments showing that the E/F suubnit (*E*-state) interface of RhoEmpty aligns almost perfectly with an open-ring intersubunit interface (left), as opposed contrast to the B/C subunit (*T^*^*^-^ state) subunit interface of Rho^PolyU-P2_1_^, which has adopted a right-handed orientation (right, red arrows).

Inspection of the E/F active site shows that the *E*-state is unique within the closed-ring Rho hexamer in adopting a relative intersubunit orientation markedly different from that of the other protomers. This altered configuration arises due to a transition of subunit F (which is bounded by subunits A and E, respectively) from the top step of the helical RNA staircase to the bottom (**Fig. 2b**). Subunit F is also the only protomer that does not contact RNA in any closed-ring structure determined so far, and its interface with subunit E in the Rho^Empty^ structure represents a completely substrate-free component of the catalytic cycle. Given these considerations, we compared the intersubunit positions of the E/F protomer conformation of the back-soaked model with those of a completely RNA-free, open-ring Rho structure of Rho determined previously. Interestingly, the intersubunit orientation of motor domains in the open-ring structure (PDB: 1PVO), which all assemble with a left-handed helical pitch, proved to be a very close match to the orientation of the E/F protomer pair in both the Rho^Empty^ and Rho^PolyU-P2_1_^ structures, but distinct from other subunit dimers in the RNA-bound models (which adopt a right-handed pitch) (**Fig 4e**). This congruency indicates that, upon ATPase product and RNA release, a single interface in the Rho hexamer relaxes to an energetically favorable conformation present in the open-ring Rho structure. Therefore, the nucleotide-dependent structural transition, which promotes ring opening and closing during RNA loading, also plays a direct role in catalytic cycling.

### Sequence and ATPase state-specific differences in RNA binding by Rho

The ATPase activity of *E. coli* Rho has been well-established to depend not just on RNA binding (59), but also on RNA sequence, with pyrimidine-rich RNAs providing a greater increase in ATP turnover compared to purine-rich substrates (37, 38). To better understand this dependency, we crystallized and determined the structure of Rho in complex with a poly-purine RNA (rA_12_) and ADP•BeF3 (hereafter referred to as Rho^PolyA^), and compared it to the other Rho structures determined thus far (**Fig. 5**). Notably, the Rho•poly-A complex co-crystallized in the same P2_1_ space group and unit cell as the new closed-ring Rho model bound to poly-U. The poly-A substrate similarly forms an overwound, single-stranded RNA helix in the Rho pore that is highly similar to that formed by poly-U (**Fig. 5 and Supplementary Fig. 7**), despite the extra bulk afforded by the additional five-member ring on the purine bases. This finding indicates that the tight coiling of RNA seen in the original Rho model is not an effect of RNA sequence or crystal packing, but likely reflects the standard manner by which the helicase encircles substrate during translocation.

**Figure 5.**
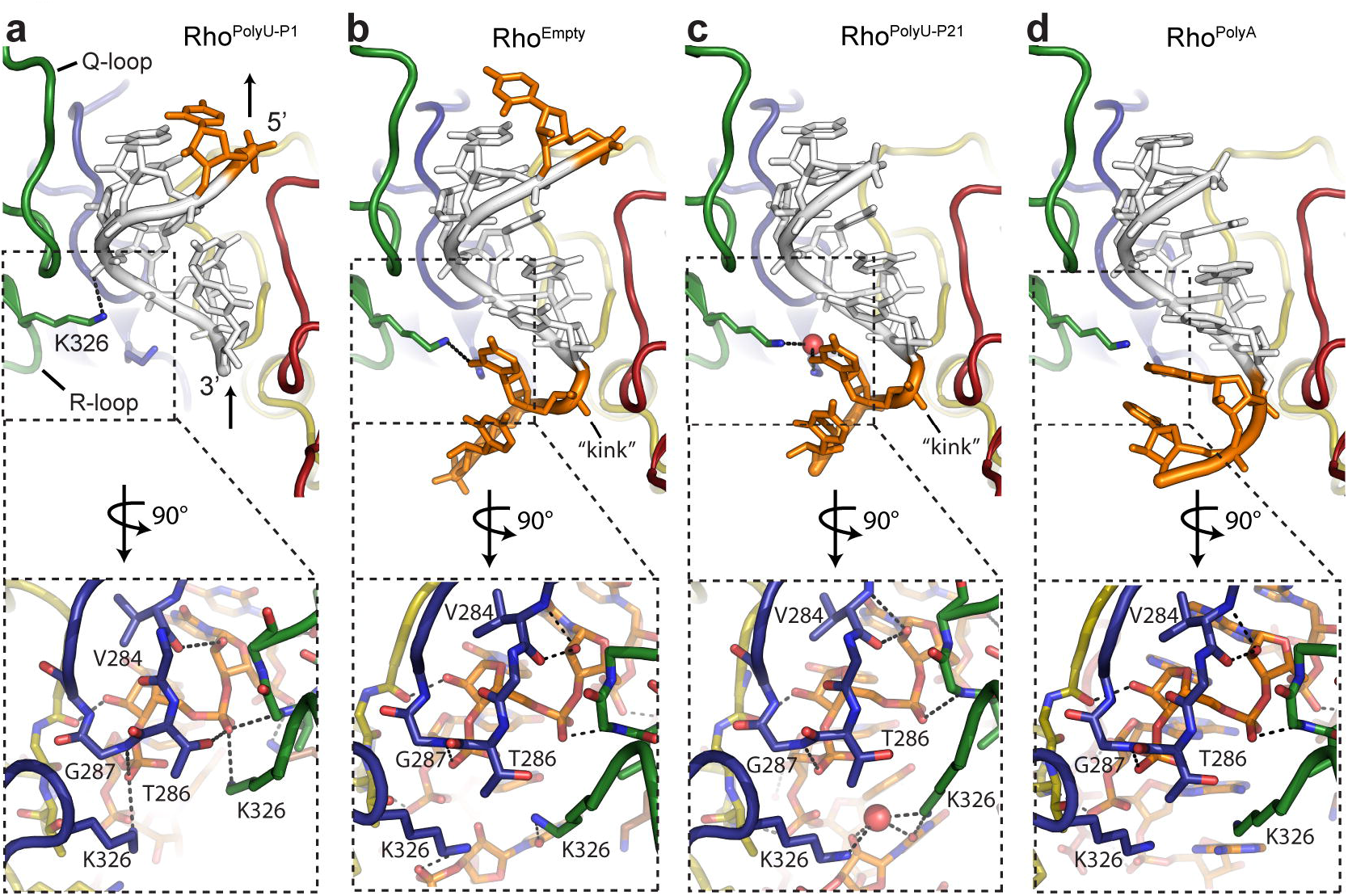
Comparison of RNA binding among closed-ring Rho structures. The five core RNA bases common to all structures are colored white, while additional 5’- and 3’-bases are colored orange. Arrows at the 5’ and 3’ end of the bound RNA indicate the direction of RNA translocation relative to the protein. Protein subunits are colored as in **Fig. 2a**. Insets reveal protein-RNA hydrogen bonding details (dashed lines) at the B/C subunit interface. (**a**) Rho^PolyU-P1^ contains six bases of RNA (orange/white) bound by the Rho Q- and R-loops in a spiral staircase conformation. (**b**) The Rho^Empty^ structure allows for the modeling of eight bases of RNA (orange/white). Compared to the Rho^PolyU-P2_1_^ structure, there is additional density for two 3’ bases in a kinked conformation. (**c**) The Rho^PolyU-P2_1_^ structure allows for modeling of seven bases of RNA (orange/white). Compared to the Rho^Empty^ structure, density is lacking for the 5’ base. This structure reveals a hydrogen bonding network stabilizing interactions between R-loop residue K326 of subunits B and C, a water molecule, and the 2’-OH and uracil base of the bound RNA kink. (**d**) The structure of Rho^PolyA^ solved under similar conditions to Rho^PolyU-P2_1_^, also reveals two additional 3’ bases, but in a pseudo A-form helical arrangement, rather than the kinked conformation; in contrast to the H-bonding network observed with the kinked PolyU substrate, no additional protein-RNA contacts are formed.

Close inspection of the poly-A Rho model with its poly-U counterparts reveals notable differences in the specifics of RNA binding. Interestingly, these differences are evident not just between purine- and pyrimidine-bound Rho hexamers, but also between the different poly-U models imaged thus far (**Fig. 5a-c**). In particular, the Rho^PolyU-P2_1_^ and Rho^PolyA^ structures lack electron density for the 5’-most base present in original Rho^PolyU-P1^ structure, and instead manifest new density for two additional bases at the 3’-end of the RNA (**Fig. 5 and Supplementary Fig. 7b-g**). However, the bound RNAs also show conformational differences between these three structures, with the additional 3’ nucleotides visible in the poly-A bound Rho model extending in a straight line from the helicase pore, but in the new poly-U structures folding back into a tight kink that deviates from the interior RNA spiral configuration but maintains base-stacking interactions (**Fig. 5 and Supplementary Fig. 7**). The U-U kink seen in the poly-U models allows for a network of contacts to be formed between the ribose 2’-OH and the uracil O2 atom of the penultimate nucleotide, a newly evident water molecule, and Lys326 of both subunits B and C (the latter of which alters the contact that had seen previously between this invariant amino acid and the backbone phosphate of RNA in the original Rho^PolyU-P1^ hexamer) (**Fig. 5a-c, Supplementary Fig. 8**). Interestingly, in analyzing the Rho^Empty^ structure, we observed electron density for the additional, kinked 3’ bases of RNA similar to that present in Rho^PolyU-P2_1_^; however, this model also exhibited density for the 5’-most base seen in Rho^PolyU-P1^ (**Fig. 5b and Supplementary Fig. 7b-d**). Thus, upon loss of nucleotide and opening of the *E*-state active site, the poly-U RNA bound to Rho^Empty^ appears to adopt a hybrid conformation between the original Rho^PolyU-P1^ crystal form and the new Rho-P2_1_ crystal form.

Concomitant with the distinct 5’- and 3’-end structures of different bound RNAs, specific protein-RNA contacts also undergo a change. Overall, the new poly-U and poly-A models form a similar number of total polar contacts between Rho and RNA when compared to the prior Rho^PolyU-P1^ structure, but these frequently involve different residues. In addition to the changes in Lys326 noted above, Thr286 of Rho’s Q-loop (the L1-loop equivalent of DnaB and T7 gp4) loses its interaction with the phosphate of RNA that was seen in the original closed-ring structure of Rho, leading both the backbone amide (not observed in RhoPolyU-P1) and carbonyl (observed in Rho^PolyU-P1^) groups of Val284 to shift and make a new contact with the 2’-OH of its associated ribose (**Fig. 5a-d**). These three models also manifest a slightly steeper RNA coil that is less compressed compared to the RNA seen previously in the P1 Rho model, likely due to changes in the intersubunit orientations between protomers E/F and F/A (**Fig. 2e and Supplementary Fig. 7**).

### A conserved allosteric helix coordinates crosstalk between subunits and their catalytic centers

In considering how changes in the ATPase centers of our closed-ring Rho models might be coupled to RNA binding, we used the higher-resolution structures determined here to look more closely at the linkages that mediate intersubunit contacts between the nucleic acid and nucleotide binding sites. By co-aligning all six pairwise interfaces of the Rho^PolyU-P2_1_^ model, using the ATP-binding P-loop as an reference point, we found that a single α-helix (Rα1), which contains residues responsible for forming the salt-bridge network linking the RNA binding pore to the ATPase center, undergoes a large range of previously-unnoticed hinge movements (**Fig. 6a and Supplementary Fig. 6**). The N-terminus of this helix (which corresponds to the “R-loop” of Rho and the “L2”-loop of RecA helicases such as DnaB and T7 gp4) binds RNA via K326 (**Fig. 5**), while the C-terminus of this helix contains backbone carbonyls that interact directly with the catalytic water molecule and the arginine-based γ-phosphate sensors in the two T^*^ states (**Figs. 3b and 6b**). This movement appears to be caused by an intersubunit levering motion of ∼10Å at the helix’s N-terminus, which alters the interactions in an ordered manner that correlates with the ATPase state of each subunit (**Fig. 6b and Supplemental Movie 1**). This observation suggests that R-loop mediated RNA binding, aided by interactions within an intersubunit allosteric network, not only positions the catalytic glutamate via the arginine switch residue as described previously (29), but also directly controls the position the arginine finger, arginine valve, and catalytic water by coupled movements with associate Rα1 helix.

**Figure 6.**
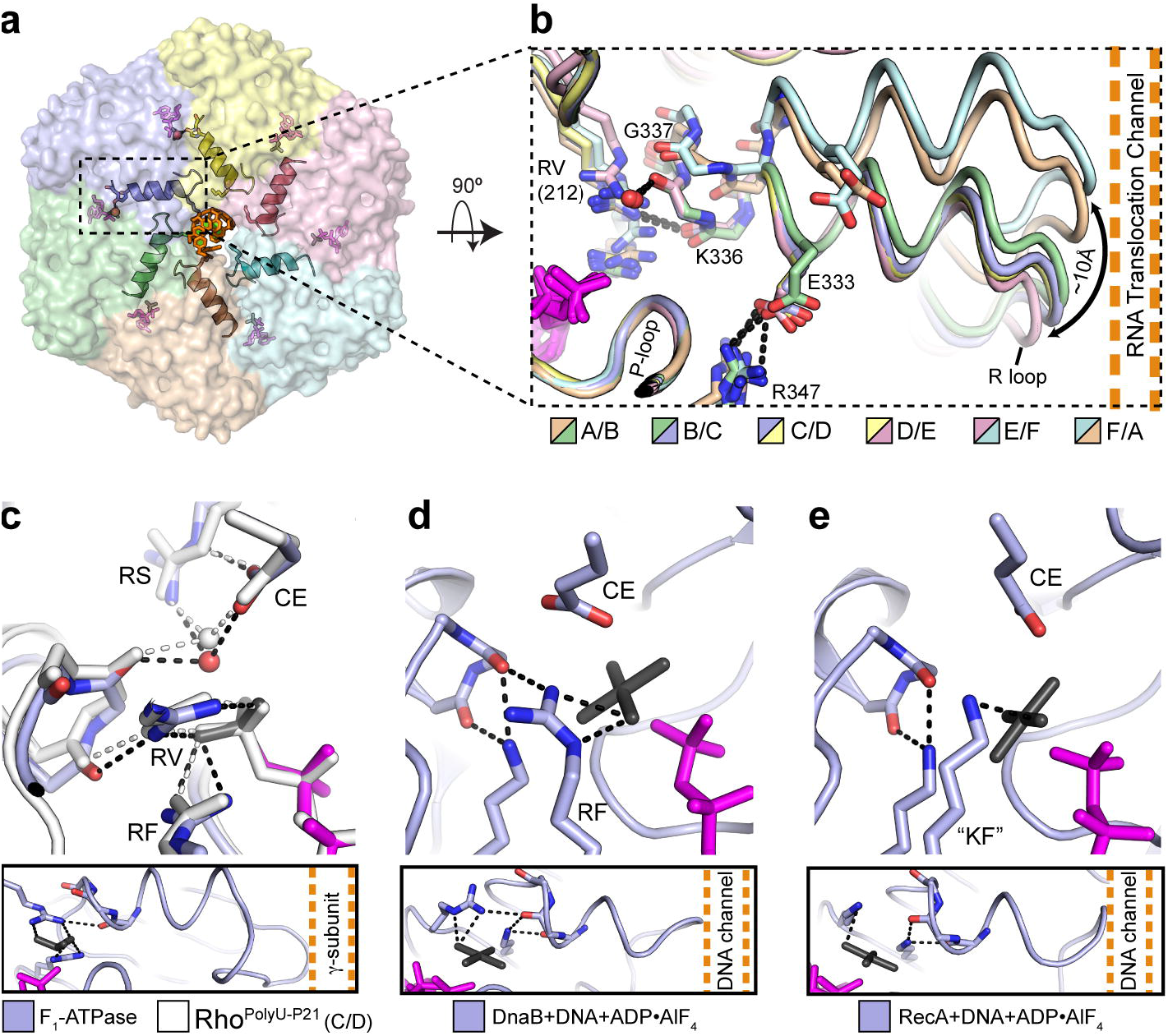
A conserved allosteric helix couples pore loop movement and ATPase status in RecA-like hexameric motors. (**a**) An α-helix (Rα1, cartoon) in each subunit of the Rho hexameric motor domain (transparent surface) directly connects the RNA and ATP binding sites. RNA – orange, nucleotide – magenta, catalytic water molecules – red spheres (subunits B and C only). Lys326 (colored sticks) directly connects the allosteric helix to bound RNA. (**b**) Alignment of all active sites in Rho^PolyU-P2_1_^ reveals the consequences of inter-subunit conformational transitions in the allosteric helix. The helix itself directly links the RNA binding R-loops in the center of the ring to the position of the γ-phosphate sensing residues in the ATP-binding sites at the periphery of the ring, as well as to the catalytic water in the *T^*^*active sites. Dashed lines (colored by interface) show bonding interactions present in the B/C, C/D and D/E subunit interfaces. Abbreviations: RV – arginine valve, RF – arginine finger. (**c**) Alignment of the catalytic site in the C/D subunit interface of Rho^PolyU-P2_1_^ (white) with the catalytic site in the structure of the F_1_-ATPase bound to ADP•BeF3 (colored, PDB ID: 1W0J) reveals nearly identical alignments of the catalytic residues and the backbone carbonyl groups of the allosteric helix (60). Lower panel – position of the F1 allosteric helix with respect to the rotating γ-subunit shaft. (**d**) The structure of DnaB bound to nucleic acid and GDP•AlF_4_ reveals interactions between its analog of helix Rα1 and γ-phosphate sensing residues (31). (**e**) The structure of RecA bound to double-stranded DNA and ADP•AlF_4_ reveals a similar helix configuration as in panels **b-d**(61). KF – lysine finger.

In conducting this analysis, we note that the α-helix corresponding to Rα1 is a conserved feature of the RecA fold, and is present not only in Rho’s closest homolog (the F1-ATPase, (60)), but also in more distantly related RecA-family hexameric helicases and translocases. In all cases, the C-terminal carbonyls of the helix equivalent to Rα1 contact amino acids that interact with the γ-phosphate moiety (and catalytic waters when present), and that appear poised to help control the active-site organization of their respective hexameric motors through a similar mechanism (**Fig. 6c-e**) (31, 60, 61). These comparisons indicate that this helix, which lies downstream of the R/L2 loop region, is likely involved in allosterically coupling substrate movements to ATPase status in RecA-like motors in general.

## DISCUSSION

### A possible role for ring distortion and strain in the mechanism of Rho and other hexameric motors

In the present study, we have used solution-based structural methods and X-ray crystallography to better understand how a RecA-family hexameric motor protein, Rho, couples changes in nucleotide status to conformational transitions that underpin ring loading and translocation events. Our SAXS data confirm that two distinct open- and closed-ring structures are accessed prior to and following RNA engagement, respectively (**Fig. 1a-c**), suggesting that the transition between these two states likely occurs as a relatively simple rigid body rotation of subunits with respect to each other (**Supplemental Movie 2**). Notably, our studies indicate that at nucleotide concentrations approaching those in the cell (values between 2 and 10 mM ATP have been measured for *E. coli* (62-64)), Rho spontaneously and preferentially occupies an open-ring state, a form that is well-suited for engaging nascent RNA transcripts. A need for contacts engendered by both substrate nucleic acid and ATP binding to cooperatively induce ring closure is consistent with the idea that formation of a translocation-competent state may capture and harness strain as a motive force to help promote nucleotide cycling. Consistent with this model, the formation of a relaxed, left-handed inter-subunit orientation characteristic of the open-ring structure is seen only in the product release state, which also releases its grip on RNA to reset the RNA binding cycle (**Fig 4e**).

Strain imparted by substrate-induced conformational changes may underlie ATP cycling in other, more distantly-related hexameric helicases. For example, recent structures of DnaB-family helicases have shown that the ATPase domains of these hexameric motors adopt an uncoupled, planar conformation in the absence of nucleic acid substrate (65), but form a steep right-handed helical conformation with a single open interface when DNA and GDP•AlF4 are present (31, 65). Electron microscopy data on the eukaryotic minichromosome maintenance proteins (MCMs) have revealed that a left-handed open-ring conformation is the default state of metazoan MCM hexamers (27), and that the Cdc45•GINS complex, along with ATP and DNA are needed to drive the formation of a stable closed ring with an internal, right-handed pitch (26, 34). Together, these studies support the idea that nucleic acids cooperatively activate ATP hydrolysis in ring-shaped many RNA and DNA translocases by serving as a template for the asymmetric assembly of ATPase subunits. This asymmetry in turn promotes the formation of a defined sequence and order of active site states (29, 30), which allows the energetics of cooperative substrate binding in one subunit to be allosterically coupled to cooperative product release in another. One exception to this trend may be the papilloma virus E1 helicase, which has been found to adopt an asymmetric, right-handed closed ring in the absence of ATP and DNA (66).

### RNA sequence dependencies of the Rho functional cycle

In addition to the insights afforded into ring dynamics and the establishment of a sequential ATPase order, the structures presented here also help to explain some of the known effects that RNA sequence has on Rho’s ATPase activity. In general, the molecular basis for substrate-specific sequence dependencies in hexameric helicases and translocases has remained relatively ill-defined. Some ring-shaped helicases, such as T7 gp4, decelerate when confronted with GC-rich duplexes (67), indicating that motor function can be impacted by substrate stability.

For Rho, transcription termination activity is stimulated by cytosine-rich sequences (13), due in part to the recognition requirements of its primary RNA (*rut*)-binding sites (10, 38), and also to the ability of primary site occupancy to promote Rho ring closure (see accompanying study (45)). However, Rho’s RNA-stimulated ATPase and translocation activities also appear to be directly affected by pyrimidine-specific sequence preferences independent of primary site interactions (36-38).

The differences in secondary site RNA binding seen here between poly-U and poly-A bound forms of Rho (along with the preferential ring closure capability of poly-U over poly-A described in the accompanying study (45)) indicate that sequence discrimination in the helicase may in part arise from interrogation of RNA sequence at the point where the nucleic acid threads into the ring. In particular, the 3’ kink seen in the new Rho^PolyU-P2_1_^ structure allows a ribose 2’- OH and two pyrimidine bases to interact with an invariant lysine, Lys326, on two different R-loops. These contacts are absent in the 3’ RNA region when Rho is bound to poly-A, and in the original Rho^PolyU-P1^ model (where the 3’ nucleotides are unstructured, see **Fig. 5**). Intriguingly, the lysines involved in binding the U-U kink in Rho^PolyU-P2_1_^ project from subunits B and C, which also manifest the two *T^*^* states observed in the hexamer (**Fig. 2a**). These interactions suggest that Lys326, in addition to its role neutralizing the backbone charge of RNA bound in Rho’s translocation pore (29), may also serve as a sensor for RNA sequence to help properly organize the B- and C-subunit active sites (**Fig. 3a**). Such a function would explain why pyrimidine-rich *rut* sites are particularly well-suited as secondary site ligands to promote nucleotide hydrolysis by Rho, in that they provide an RNA structural motif that helps set up an idealized ATPase configuration.

The differences in the 3’ RNA structure seen here for the purine and pyrimidine substrates likewise have implications for understanding how RNA sequence is coupled to step size. For instance, using nucleotide analog interference mapping (NAIM) to insert dNTPs into RNA, Rho has been found to be inhibited by the presence of deoxyribose at intervals of every seven nucleotides, with the level of inhibition dependent on the local sequence environment (36). Our structures reveal that pyrimidine, but not purine substrates can specifically interact with Lys236 of Rho by forming a specific kink that involves the 2’OH and pyrimidine base of the incoming 3’-RNA end (**Fig. 5**). Given that movements of the R-loop are directly coupled to changes in the allosteric network that couples ATPase status with RNA-binding (**Figs. 6 and Supplementary Fig. 6**) (29), the ability to adopt a kink by pyrimidines may help underpin the positional and sequence dependency of the 2’-OH activation step seen to occur roughly every seven steps by NAIM (36, 68).

### Structural snapshots reveal translocation sub-steps in a hexameric motor protein

The present study provides the most comprehensive structural analysis to date for a hexameric helicase bound to different nucleic acid substrates and nucleotide intermediates. In comparing the original P1 and the new P2_1_poly-U bound models of Rho, both hexamers can be seen to possess a single *E*-state active site (subunit E) that sits adjacent to a *T*(ATP-bound) state ATPase center (subunit D). The two structures diverge, however, in that the Rho^PolyU-P1^ form contains a second *T*-state active site (subunit C) followed by two hydrolysis-like *T^*^* states (subunits B and A), whereas the Rho^PolyU-P2_1_^ model is shifted, possessing a lone *T*-state followed immediately by two *T^*^* catalytic centers (**Fig. 2**). As a consequence, the Rho^PolyU-P1^ structure contains only a single product (*D*) state (subunit F), whereas the corresponding Rho^PolyU-P2_1_^ structure contains two (subunits F and A). This observation is consistent with prior studies showing that there exists a 2-4 subunit delay in ATP hydrolysis and product release in Rho and related RecA-family hexameric helicases such as T7 gp4 (see (69) and **Supplementary Discussion**).

Although the *E*-state active site (subunit E) remains the most solvent-accessible ATPase center in both crystal forms of Rho, changes in the relative disposition of key functional amino acids are nonetheless evident for this region between the P1 and P21 structures. In particular, the subunit E active site of the new Rho^PolyU-P2_1_^ model exhibits a dramatically displaced and off-line catalytic glutamate, but forms a new contact between the arginine-finger and the BeF3 moiety; by contrast, in the original Rho^PolyU-P1^ structure, the correct positioning of the catalytic glutamate is maintained but the arginine-finger/nucleotide contact is broken (**Fig. 3d-e**). Concomitant with these changes, two bases at the 3’ end of the bound RNA become ordered and coordinated in the RhoPolyU-P2^1^ structure, whereas in the Rho^PolyU-P1^ model, only a single extra base is apparent at the 5’ RNA end (**Fig. 5**). Interestingly, back-soaking of the new poly-U Rho crystal form, which results in nucleotide dissociation from the E-subunit interface, produces a hybrid state in which RNA density is evident at both 5’ and 3’ ends.

Taken together, the suite of Rho structures not only recapitulates the movements expected for 5’→3’ translocation along RNA, but also highlights the existence of sub-steps within the context of a sequential ATPase mechanism (**Figure 7 and Supplemental Movie 3**). In this scheme, formation of an empty ATPase center would promote downstream (3’) contacts between Rho and its substrate, while maintaining 5’ interactions leftover from the prior round of hydrolysis. ATP binding would next disrupt the 5’ RNA contact, initiating a new translocation cycle to pull the 3’ end of the bound RNA into the motor pore. Finally, product formation would complete the translocation step, helping to shepherd the 5’ RNA end out of the helicase, after which ADP release would reset the system by forming new contacts to the incoming 3’ base.

**Figure 7.**
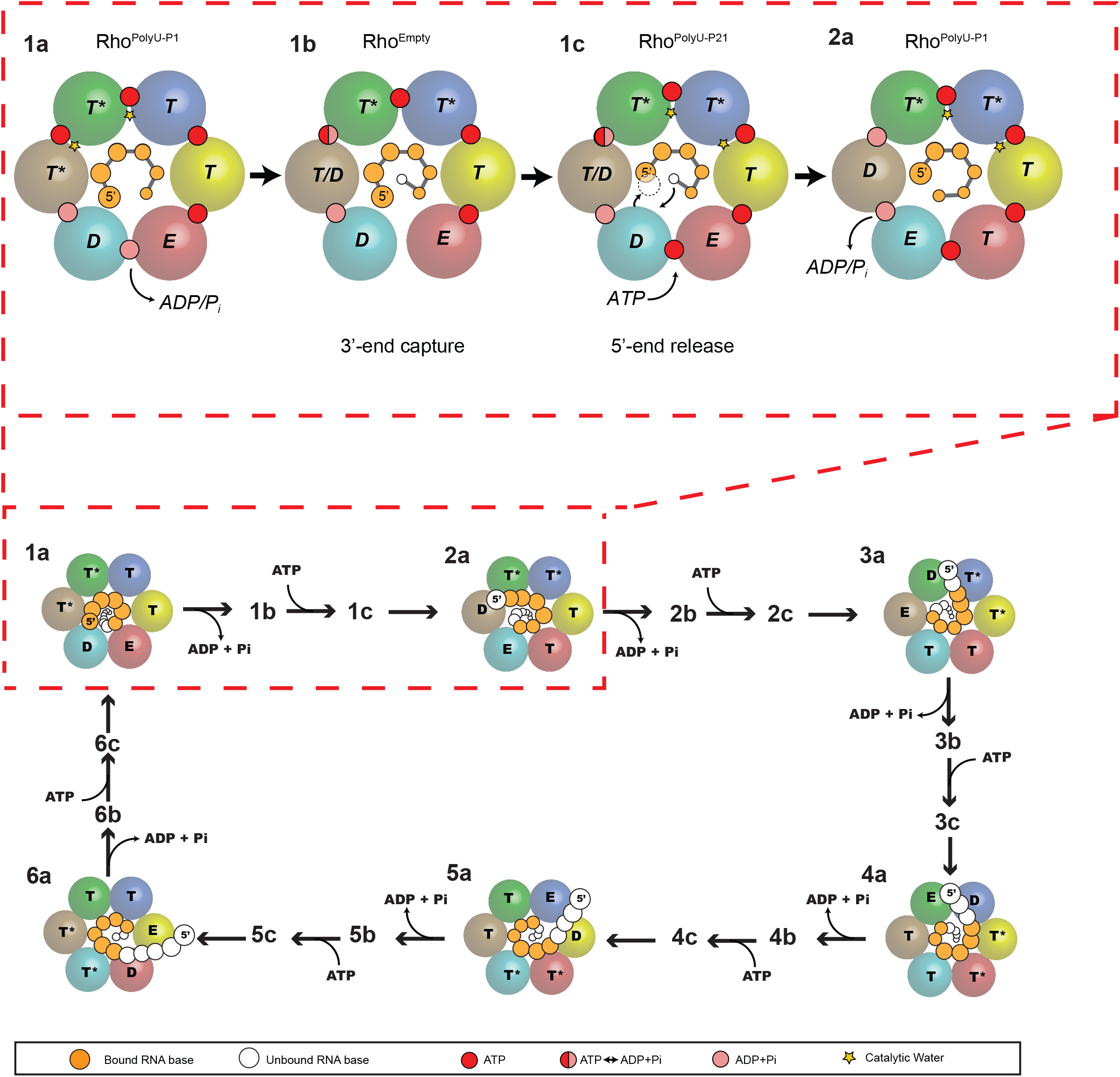
Sub-stepping in a hexameric helicase. (*Top, red dashed box*) Diagram of RNA bases and catalytic states in the Rho^PolyU-P1^, Rho^PolyU-P2_1_^ and Rho^Empty^ structures. Protein subunits are shown as large spheres colored by chain as in **Fig. 2a**. Arrows indicate the sequence of proposed sub-steps in which an incoming 3’ RNA base is bound and a 5’ RNA base is released consistent release of nucleotide and a shift in position of the *T^*^*catalytic states. Yellow stars designate the sites of the nucleophilic water observed in the various Rho structures; only in the Rho^PolyU-P2_1_^ structure does this water adopt an in-line attack position. (*Bottom*) Relationship of the substeps to a complete series of six sequential hydrolysis and translocation steps by Rho.

Although conformational substeps have been observed structurally for superfamily 1 and 2 helicases and for motors such as the F1 ATPase, myosin, and kinesin (70-73), similar insights have not been forthcoming for hexameric helicase/translocase systems. Functional substeps (as well as macrosteps) are frequently observed in single molecule studies of motor proteins (74, 75), but the architectural underpinnings of these events often remained unresolved. The present work demonstrates the existence of distinct physical substates in a RecA-family hexameric helicase, and moreover implicates both RNA binding and different stages of the ATPase cycle in the formation and dissolution of these states. Future efforts involving high-resolution single-molecule studies and structure-guided mutagenesis will help establish the timing by which different states interconvert with each other and the role that specific, protein-mediated contacts between RNA and ATP play in controlling these transitions.

## METHODS

### Small Angle X-ray Scattering (SAXS) data collection and analysis

Native Rho protein was prepared in a similar manner to selenomethionine-labeled protein as described (22). Rho protein (40 mg/mL) in size exclusion buffer (50 mM Tris 7.5, 500 mM KCl, 10% Glycerol, 1mM TCEP) was dialyzed overnight into SAXS buffer (150 mM KCl, 50 mM HEPES pH 7.5, 5 mM MgCl_2_, 5% Glycerol). Dialyzed protein was then further diluted to 5 mg/mL in SAXS buffer containing various concentrations of RNA (rU12) or ADP•BeF_3_, aliquoted into a 96 well plate, and used for data collection at room temperature using an automated system at ALS Beamline 12.3.1. Raw SAXS data were integrated, scaled, and buffer subtracted with the program OGRE (Greg Hura, Lawrence Berkeley National Laboratory). PRIMUS was used to merge data from a one second and ten second exposure and to manually remove outliers near the beam-stop in order to obtain high quality scattering curves with very little noise (76). Guinier analysis was performed using AUTORG (77). Distance distribution functions (P(r)), maximum diameters (D_max_) and real- and reciprocal-space radius of gyration (R_g_) values were calculated with GNOM (78). Structural models were generated using an open-ring Rho model (PDB: 1PVO (22)) and the closed ring crystal structure determined here (Rho^PolyU-P2_1_^). Missing loops or domains were added to the structural models by superposition with other structures or subunits containing the missing residues. OLIGOMER and FOXS were used to compare theoretical scattering curves to the data and compute the relative contributions of multi-component datasets (42, 43, 79). Background subtraction as implemented in FOXS or OLIGOMER was used to account for errors in buffer composition and improve the fit of the high-resolution data.

### Protein crystallization and data collection

Selenomethionine labeled *E. coli* Rho protein was prepared, and the RNA-ADP•BeF3 complexes were formed as described previously (22, 29), except that Rho was dialyzed into a buffer containing 10 mM Tris pH 7.5, 100 mM NaCl and 0.5 mM TCEP prior to complex formation. RNA (rU12 and rA12 polymers) was purchased from IDT. Rho was crystallized by mixing 250 nL of Rho complex (20 mg/mL) with 250 nL of a well solution containing 200 mM KOAc, 40% MPD and 0.5% ethyl acetate using a Mosquito Nanoliter Pipetting system (TTP Biotech), in a hanging-drop vapor diffusion format. Crystals grew within three days and reached maximum size within one week. Crystals were harvested directly by looping and plunging in liquid nitrogen. Crystals were mounted on bendable cryo-loops (Hampton Research), oriented to minimize spot overlap, and used for collecting diffraction data on Beamline 8.3.1 at the Advanced Light Source (80). Diffraction data from the new Rho^PolyU-P2_1_^ crystal form were significantly improved compared to the previous Rho^PolyU-P1^model (29), displaying lower mosaicity (∼1^°^ *versus* 2^°)^, and higher resolution (2.6 Å *versus* 2.8 Å). Co-crystals of Rho with a short poly-adenosine RNA (rA12) and ADP•BeF3 were obtained under nearly identical conditions by substituting rA12 RNA for rU12 RNA during rho complex formation. To obtain the Rho^Empty^ crystals, we first grew Rho^PolyU-P2_1_^ crystals, and then conducted stepwise exchange of the mother liquor with an identical solution containing no ADP•BeF3. After a series of 2-fold dilutions and equilibrations conducted over ∼15 minutes, the concentration of ADP•BeF3 in the mother liquor was reduced approximately 30-fold (further dilution caused the crystals to dissolve).

### Structure solution and refinement

All data were processed in HKL-2000 (81). Molecular replacement/ single wavelength anomalous dispersion (MR/SAD) methods as implemented in PHENIX were used to phase the Rho^PolyU-P2_1_^ structure (82). Molecular replacement was conducted using a monomer from the Rho^PolyU-P1^ structure. Phases from the resulting solution were used by the program PHASER to locate 92 selenium sites in the asymmetric unit and phase the structure using SAD (83), producing a figure of merit (FOM) of 0.39 for unmodified phases. Multi-domain, six-fold non-crystallographic symmetry (NCS) averaging and phase extension into an isomorphous, low-energy remote dataset using RESOLVE produced a FOM of 0.63 (84). These high quality maps allowed us to clearly locate bound ligands (**Fig. 2c-d and Supplementary Fig. 6a-i**) and correct small errors in previous structures (**Supplementary Fig. 3**). The final model, which was refined to an R_work/R_R_free_ of 21.8/24.6%, currently represents the highest resolution view of a hexameric helicase bound to both nucleic acid and an ATP mimic. The Ramachandran statistics for this model are 98.17% favored, 1.79% allowed, and 0.04% outliers as reported by MOLPROBITY (85).

The nearly isomorphous Rho^Empty^ and Rho^PolyA^ structures were solved by molecular replacement, using the entire Rho^PolyU-P2_1_^ structure as a search model, and R_free_ was calculated using the same test set for all structures. Rigid body refinement of the N (1-126) and C (127-417) terminal domain of each individual subunit was conducted following molecular replacement for all structures. The structures were refined in PHENIX using custom bond and angle restraints for the bound ADP•BeF3 complex, individual (Rho^PolyU-P2_1_^) or group (Rho^PolyA^ and Rho^Empty^) B-factor modeling, and TLS modeling using selections obtained from the TLSMD server (82, 86). The Rho^PolyA^ structure was refined to 3.15 Å with an R_work/R__free_ of 23.3/26.5%, contains six bound ADP•BeF_3_•Mg^2%^ moieties and resolves seven bases of poly-adenosine RNA (**Table 1 and Supplementary Figure 5f-g**). The Ramachandran statistics for this structure are 97.48% favored, 2.52% allowed, and 0% outliers. The RhoPolyA model closely aligns with both the Rho^PolyU-P1^ and Rho^PolyU-P2_1_^ structures (RMSDs of 1.57 Å and 0.82 Å respectively for backbone C-α atoms). The Rho^Empty^ structure was refined to 3.2 Å resolution with an Rwork/Rfree of 23.2/26.6%, contains five clearly resolved ADP•BeF3•Mg2+ complexes and resolves eight bases of poly-U RNA (**Table 1 and Supplementary Figure 5h-i**). The Ramachandran statistics for this structure are 97.68% preferred, 2.32% allowed, and 0% outliers. The structure displays the largest changes in overall architecture compared to the other Rho states, aligning with Rho^PolyU-P1^, Rho^PolyU-P2_1_^ and Rho^PolyA^ structures with Cα RMSDs of 2.39 Å, 0.82 Å, and 0.88 Å respectively.

### Structural analysis

All figures were prepared using PYMOL (Schrödinger). Structural superpositions were conducted with the CEALIGN PYMOL plugin (87). Energy minimized linear interpolations were computed using a CNS script written by the Yale Morph Server (88). To place atomic B-factors from different structures on the same relative scale (**Supplementary Figure 4**), atomic B-factors for each structure were divided by the average B-factor for that structure and then multiplied by the average B-factor for all structures. Distance vectors calculated between various Rho structures were calculated as described (89). Vector length corresponds to distances between Cα atoms among different structures multiplied by a scale factor of three to assist visualization, while vector color corresponds to relative distance (dark blue: 0 Å; red: 1.5 Å or greater).

### Accession codes

Crystallographic data and structural models for Rho^PolyU-P2_1_^, Rho^PolyA^ and Rho^Empty^ have been deposited into the Protein Data Bank under accession codes 5JJI, 5JJK, and 5JJL respectively.

## ACKNOWLEDGEMENTS

The authors thank Berger lab members for helpful discussions, Greg Hura and Dina Schneidman-Duhovny for guidance on SAXS data processing, Jamie Cate for assistance with distance vector calculations, Jane Tanamachi, George Meigs and James Holton at ALS Beamline 8.3.1, and Michal Hammel at ALS Beamline 12.3.1 for assistance with SAXS data collection. This research was supported by funding from the G. Harold and Leila Y. Mathers Foundation and the NIH (GM071747).

## AUTHOR CONTRIBUTIONS

NDT and JMB designed the study; QS identified the new Rho P2 _1_ crystallization condition; LBW performed protein purification, crystallization, and back-soaking to obtain the Rho^PolyA^ and Rho^Empty^ crystals; NDT performed all other protein purification and crystallization experiments, prepared SAXS samples, collected and analyzed all X-ray data, and solved and refined all structures. NDT, MRL, and JMB wrote the paper.

